# Arabidopsis *SUFB* and CLP protease regulate SUFBC_2_D to manipulate iron-sulfur cluster biosynthesis in chloroplast

**DOI:** 10.1101/2024.11.01.621566

**Authors:** Yuting Cheng, Zhaoyang Liu, Bing Yang, Qingsong Jiao, Hisashi Ito, Atsushi Takabayashi, Ryouichi Tanaka, Ting Jia, Xueyun Hu

## Abstract

Iron‒sulfur (Fe‒S) clusters are essential cofactors for Fe‒S proteins. SUFBC_2_D complex is the scaffold responsible for Fe‒S cluster assembly in chloroplasts. However, the regulatory mechanism on SUFBC_2_D remains elusive. In this study, we report that the transcription of *SUFB* responds rapidly to leaf senescence, whereas the transcription of *SUFC* and *SUFD* does not. Intriguingly, their protein contents remain stable during leaf senescence. We further found that leaf death was occurred only when *SUFB* RNAi was induced, and SUFB and SUFC contents decreased much faster in the *SUFB*-RNAi lines than in the *SUFC*-RNAi lines, indicating that SUFB had a faster turnover rate than SUFC. Moreover, overexpressing *SUFB* increased the contents of SUFC and SUFD, and SUFBC_2_D, whereas overexpressing *SUFC* did not increase SUFB and SUFD. Our findings reveal that SUFB stabilizes SUFC and SUFD via forming SUFBC_2_D, whereas SUFC lacks this function. Furthermore, *SUFB* expression was sharply downregulated when the plants were subjected to iron deficiency, whereas *SUFC* and *SUFD* expression was not. Interestingly, the contents of all three SUF members decreased, indicating that plants degrade SUFBC_2_D in response to iron deficiency by downregulating *SUFB* transcription. We subsequently studied the degradation mechanism of SUFBC_2_D. Our results indicated that all SUFs are substrates of the caseinolytic protease (CLP) because they all accumulated in the CLP impaired mutant, and they physically interact with CLPS1, the substrate recognition adaptor of CLP. Collectively, our findings provide novel insights into how plants regulate SUFBC_2_D complex via *SUFB* to adapt to leaf senescence and iron deficiency.

## Introduction

Iron‒sulfur (Fe‒S) clusters are ancient and ubiquitous protein cofactors that function in catalysis, electron transport and the sensing of ambient conditions in organisms (Lill 2009). The chloroplast contains six Fe‒S cluster types: classic [2Fe‒2S], NEET-type[2Fe‒2S], Rieske-type [2Fe‒2S], [3Fe‒4S], [4Fe‒4S], and siroheme [4Fe‒ 4S] (Lu 2018). These Fe‒S clusters are employed by more than 40 chloroplast proteins that play diverse roles in numerous fundamental metabolic and regulatory pathways, such as photosynthesis, chlorophyll metabolism, reactive oxygen species (ROS) scavenging, photorespiration and nitrogen assimilation (Ravet and Pilon 2013; Mettert and Kiley 2015; Przybyla-Toscano et al. 2018).

Fe‒S cluster assembly is not a spontaneous process *in vivo*. Plastids inherit and modify the Fe-S cluster assembly machine sulfur utilization factor (SUF) system from their ancestor cyanobacteria for Fe‒S cluster assembly and transport (Ye et al. 2006). This process involves four steps: iron uptake, sulfur mobilization, Fe‒S cluster assembly and the transport of Fe‒S clusters to downstream apoproteins (Yang et al. 2023). In short, two highly similar DNAJ proteins, DJA5 and DJA6, bind iron through their cysteine residues and deliver iron to the iron recipient SUFD (Saini et al. 2010; Zhang et al. 2021). Cysteine desulfurase (NFS2) removes sulfur from free cysteine and delivers it to sulfur transferase (SUFE1), which then transfers it to SUFB in the SUFBC_2_D complex (Layer et al. 2007), where the sulfur assembles with iron (Balk and Lobréaux 2005; Hu et al. 2017). The assembled Fe‒S clusters are then delivered to downstream apoproteins, which requires the involvement of several Fe‒S cluster carriers, including NUFs (NifU-domain proteins), SUFA, high chlorophyll fluorescence 101 (HCF101), and glutaredoxins (Grxs) (Nath et al. 2016; Touraine et al. 2019). Fe-S clusters and Fe-S proteins originated under highly reducing conditions during early evolution and are sensitive to damage by ROS (Imlay 2006; Garcia et al. 2022).

In bacteria, the expression of *sufABCDSE* operon is induced under oxidative stress and iron starvation conditions (Outten et al. 2004; Lee et al. 2008), which suggests that the *sufABCDSE* operon is tightly regulated to provide Fe‒S clusters under in certain environmental conditions. In chloroplasts, oxidative stress occurs because of the production of ROS during photosynthesis; therefore, the SUF system is the sole machine used to assemble Fe‒S clusters. However, *SUFB* genes in the leaves of Arabidopsis (*Arabidopsis thaliana*) and rice (*Oryza sativa*) are downregulated in response to iron-limiting conditions (Xu et al. 2005; Liang et al. 2014). Furthermore, the Bio-Array Resource Database (http://bbc.botany.utoronto.ca/) indicates that the mRNA level of *AtNAP1* is significantly elevated in senescent leaves when compared with green leaves (Nagane et al. 2010). SUFB forms a stable complex with SUFC and SUFD with a 1:2:1 stoichiometry, as indicated by electrospray ionization‒mass spectrometry analysis and biochemical methods (Wollers et al. 2010; Hu et al. 2017). The SUFBC_2_D complex serves as the Fe‒S cluster scaffold and is conserved from bacteria to higher plants (Balk and Lobréaux 2005; Hu et al. 2017); therefore, the biogenesis of Fe–S clusters within chloroplasts may be tightly regulated through SUFBC_2_D. To date, little is known about how SUFBC_2_D is regulated in plants.

In this study, we confirmed that the transcription of *SUFB* can rapidly respond to leaf senescence and iron deficiency in plants, which is important for retaining the SUFBC_2_D content during leaf senescence and decreasing the SUFBC_2_D content under iron starvation. This is because SUFB has a faster turnover rate than SUFC. Subsequently, 2D BN/SDS‒PAGE analysis revealed that SUFB not only stabilizes SUFC and SUFD but also mediates their assembly into SUFBC_2_D, a function that SUFC lacks. Additionally, we found that SUFs all serve as degradation substrates for the CLP system. On the basis of our findings, we propose that the expression of *SUFB* and the degradation of SUFB respond to various plant statuses to regulate the content of the SUFBC_2_D complex, which is an indispensable strategy for plants to adapt to these conditions. Furthermore, we demonstrated that the CLP protease is responsible for the degradation of SUFB, SUFC and SUFD. Taken together, the results of this study provide new insights into the regulatory mechanism of SUFBC_2_D, the scaffold complex for Fe–S cluster biogenesis in plastids.

## Results

### Changes in the *SUFB* gene and SUFB protein in response to leaf senescence

It was previously reported that SUFBC_2_D is an essential scaffold complex for Fe–S cluster biosynthesis within chloroplasts (Hu et al. 2017). Nagane et al. (2010) reported markedly elevated transcript level of the *SUFB* gene during leaf senescence, whereas the transcription levels of *SUFC* and *SUFD* remained stable throughout the senescence (Nagane et al. 2010). Additionally, mRNA expression atlas data from the eFP Browser (http://bar.utoronto.ca/; Schmid et al., 2005; Winter et al., 2007) revealed that *SUFB, SUFC* and *SUFD* have different expression patterns during leaf development, and only *SUFB* is highly expressed in senescent leaves (Supplementary Fig. S1). To confirm these results, wild-type (WT) mature leaves were collected at weekly intervals from the 4- to 8-week-old plants for the analysis of *SUFB*, *SUFC*, and *SUFD* transcript levels. qRT–PCR analysis revealed that the transcript level of *SUFB* increased with age, whereas the transcript levels of *SUFC* and *SUFD* remained relatively stable during this period (Fig. 1A). This finding is consistent with those of previous studies.

**Figure 1.**
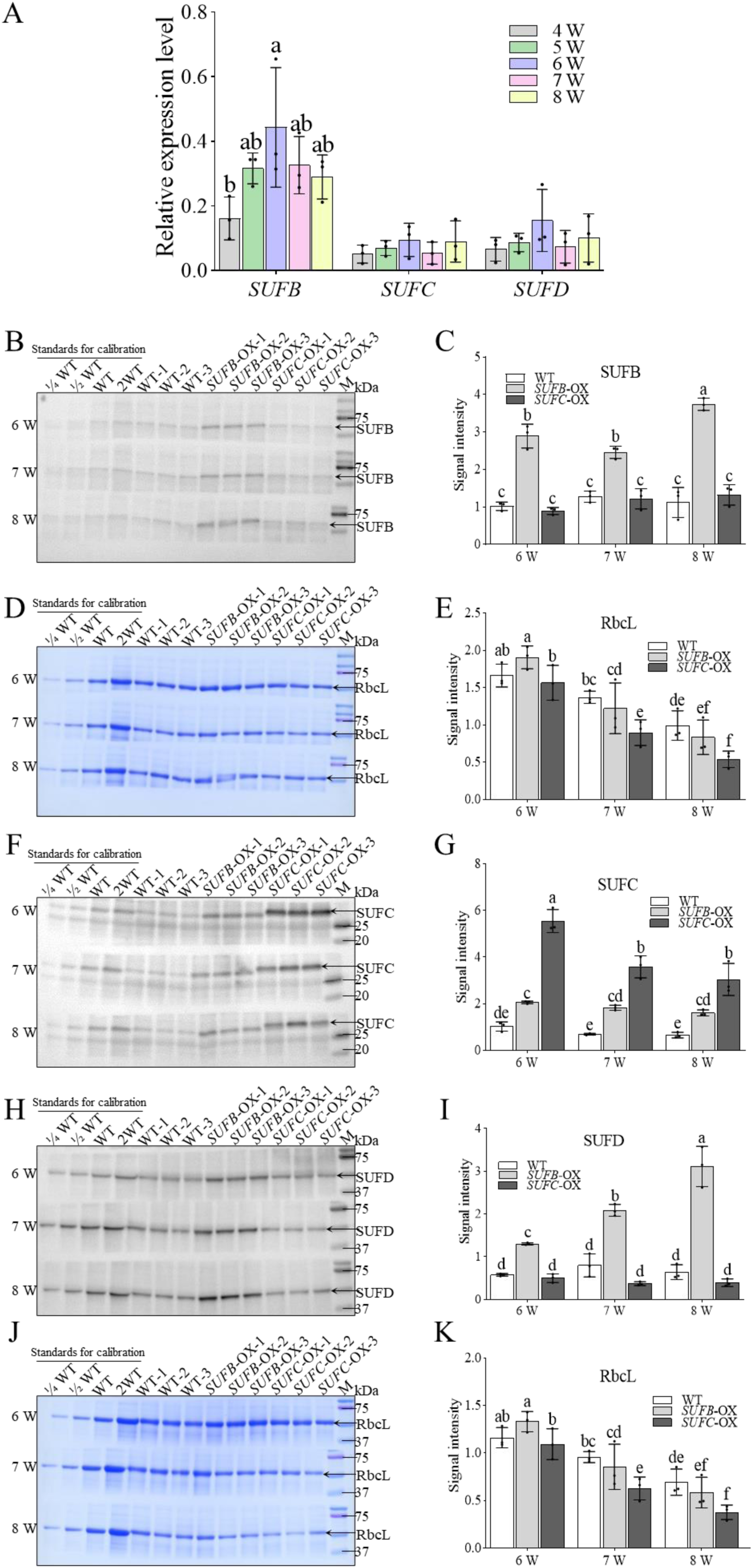
Analysis of gene expression and protein abundance during the processes of natural leaf senescence. (A) Relative expression levels of the *SUFB*, *SUFC* and *SUFD* genes in the mature leaves of 4-8-week-old WT plants. W represents the number of weeks. *ACTIN2* was used as an internal reference for normalization. The data are transcript levels relative to *ACTIN2* (with *ACTIN2* set to 1). (B, F, H) Immunoblot analysis of SUFB, SUFC and SUFD contents in the mature leaves of 6-8-week-old plants. (D, J) The corresponding Coomassie Brilliant Blue (CBB)-stained bands corresponding to the Rubisco large subunit (RbcL) are shown as loading controls in Panel B, and Panels F and H, respectively. (C, E, G, I, K) Quantification analysis of SUFB, SUFC, SUFD, and RbcL contents in 6-8-week-old plants, corresponding to Panels B, D, F, H, J on the left. The signal intensity values corresponding to the WT protein bands were defined as 1.0 and used as standards for calibration on each PVDF membrane. In Panels A, C, E, G, I, and K, the data represent the means ± SEs (n = 3 biological replicates), and different lowercase letters denote significant differences (*P* < 0.05), as determined by two-way ANOVA followed by Tukey’s post hoc test.

Having identified the differential transcriptional responses of *SUFB, SUFC*, and *SUFD* to leaf senescence, we subsequently examined the protein levels of SUFB, SUFC, and SUFD during leaf senescence by immunoblotting (Fig. 1B-K; corresponding standard curves are shown in Supplementary Fig. S2). We first focused on their content in WT (the open bars at the left in each group). Notably, the level of SUFB did not increase despite its elevated transcript level. The protein content of SUFC and SUFD also remained stable over extended periods of senescence, whereas CBB staining results indicated that the content of the Rubisco large subunit (RbcL) gradually decreased with increasing leaf age. These results suggest that the SUFBC_2_D complex must persist longer to meet the need for Fe–S clusters during leaf senescence.

The importance of Fe–S cluster formation during leaf senescence is illustrated by the fact that key chlorophyll-degrading enzymes that are crucial for this process, such as 7-hydroxymethyl chlorophyll *a* reductase (HCAR) and pheophorbide *a* oxygenase (PAO) contain Fe–S clusters (Hirashima et al. 2009; Meguro et al. 2011). The transcript level of *SUFB* increases during leaf senescence, whereas its protein level remains constant. This apparent discrepancy may be explained by the highly oxidative environment in senescent cells, which is likely to accelerate SUFB degradation. To maintain the homeostasis of the SUFBC_2_D complex, plants appear to increase *SUFB* transcription to compensate for the increased degradation rate. Our data suggest that plants maintain the crucial role of the SUFBC_2_D complex throughout the early stage of the senescence process by increasing *SUFB* transcription levels, thereby ensuring that sufficient Fe–S clusters are provided for various cellular processes at this stage.

### Decreasing SUFB content cause abnormal leaf senescence

To test the above hypothesis, the WT and inducible *SUFs*-RNAi lines were employed to analyse the leaf senescence phenotypes. On the 12th day after the inducer dexamethasone (Dex) was sprayed, the mature leaves of *SUFB*-RNAi plants were necrosis, characterized by wilting but retaining a green colour (Fig. 2A). In contrast, the corresponding leaves of the WT, *SUFC*-RNAi and *SUFD*-RNAi plants presented a normal senescence phenotype (Fig. 2A). These data strongly suggest that the transcript level of *SUFB* is more important than those of *SUFC* and *SUFD* for programmed cell death.

**Figure 2.**
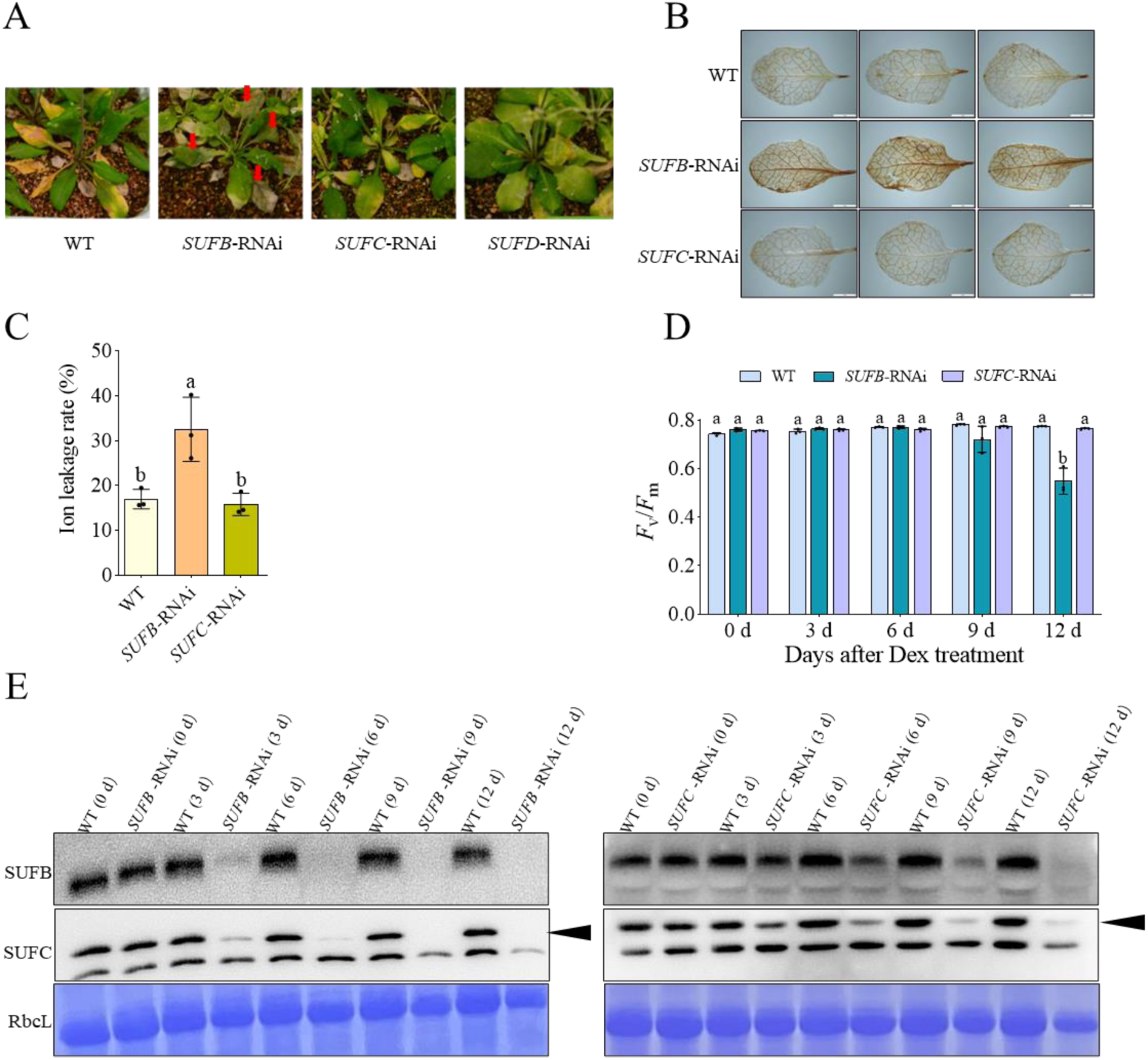
Phenotypes and biochemical analysis of the mature leaves after the induction of S*UFB, SUFC* and *SUFD* silencing. (A) Phenotypes of WT, *SUFB-*, *SUFC-* and *SUFD-*RNAi plants on the 12th day after 20 µM Dex solution was sprayed. Dex spraying was begun when the plants were 4 weeks old. The red arrows indicate that non-programmed cell death occurred in the leaves. (B) DAB staining to detect H_2_O_2_ accumulation in the leaves of WT and RNAi plants. Three-week-old plants were sprayed with 20 µM Dex once every three days. At 12th day, the same age leaves were harvested from these plants for DAB staining. Scale bars = 1 mm. (C) Ion leakage rates of the leaves of WT and RNAi plants. The leaves used for measuring electrolyte leakage were the same age and underwent the same treatment as those in (B). The data represent the means ± SEs (n = 3 biological replicates). Different lowercase letters denote significant differences (*P* < 0.05), as determined by one-way ANOVA followed by Tukey’s post hoc test. (D) *F*_v_*/F*_m_ values of WT and RNAi plants. Three-week-old plants were sprayed with 20 µM Dex once every three days, and the *F*_v_*/F*_m_ values of leaves of the same age were measured before each spraying. The data represent the means ± SEs (n = 3 biological replicates). Different lowercase letters denote significant differences (*P* < 0.05), as determined by two-way ANOVA followed by Tukey’s post hoc test. (E) Immunoblot analysis of SUFB and SUFC contents in the same WT and RNAi plant leaves whose *F_v_/F_m_* values were measured. The black arrows indicate the bands of SUFC. The smaller bands in the same blot may be the degradation products of SUFC. CBB-stained bands of the Rubisco large subunit (RbcL) are shown as loading controls.

3,3′- Diaminobenzidine (DAB) staining produced a dark brown colour in the *SUFB*-RNAi leaves, whereas a light brown colour was observed in the WT and *SUFC*-RNAi leaves (Fig. 2B), indicating that H_2_O_2_ accumulated more in the *SUFB*-RNAi leaves than in the leaves of the other lines. Additionally, we monitored the alteration in the ion leakage rate, which was significantly greater in *SUFB*-RNAi leaves than in the leaves of WT and *SUFC*-RNAi plants (Fig. 2C), indicating that suddenly inducing a decrease in SUFB biosynthesis in mature leaves caused ROS accumulation, photosystem destruction and necrosis. At the initial stage of *SUFB*- or *SUFC*-RNAi induction, the maximum quantum yield of photosystem II (PSII) photochemistry (*F*_v_/*F*_m_) values was similar to that of the WT. Up to the12th day after RNAi induction, the *F*_v_/*F*_m_ value of the mature leaves of *SUFB*-RNAi plants significantly decreased, whereas the *F*_v_/*F*_m_ values of the other lines did not change (Fig. 2D). These results suggest that SUFB plays a significant role in the senescence process and that a deficiency in SUFB biosynthesis causes oxidative damage and an accompanying decline in photosynthetic activity.

### The turnover rate of SUFB is faster than that of SUFC

To investigate why the necrosis of mature leaves occurred only when *SUFB* transcription was suppressed, we examined the changes in SUFB, and SUFC levels following the induction of *SUFB*-RNAi or *SUFC*-RNAi (Fig. 2E). Three days after the *SUFB*-RNAi plants were sprayed with the inducer, the abundance of SUFB had decreased to a very low level. On the 6th day, the content of SUFB was decreased further and was close to the limit of detection. The decrease in the SUFC content was consistent with that of the SUFB. In the mature leaves of the *SUFC*-RNAi plants, however, approximately half of SUFB and SUFC both remained at the 6th day after the first spraying of the inducer, with a further reduction to the limit of detection at the12th day (Fig. 2E). Our previous studies revealed that when RNAi was induced by 10 µM Dex, only a small amount of SUFB or SUFC was detected in newly growing leaves (Hu et al. 2017), indicating that SUFB and SUFC biosynthesis was almost completely eliminated in the *SUFB*- and *SUFC*-RNAi lines, respectively. The fact that both SUFB and SUFC decreased faster in the mature leaves of the *SUFB*-RNAi lines than in those of the *SUFC*-RNAi lines after the inducer was sprayed suggested that the SUFB turnover rate is faster than the SUFC rate when the plants are in the same complex. Both the SUFB and SUFC proteins are essential for the formation and maintenance of the SUFBC_2_D complex. The absence of either protein leads to destabilization of the entire complex and subsequent degradation of other subunits. When *SUFB* transcription is inhibited, preexisting SUFB protein is rapidly degraded. In contrast, when SUFC transcription is inhibited, the degradation of existing SUFC protein occurs over a longer timeframe. These findings suggest that the faster turnover rate of the SUFB protein is the limiting factor in maintaining the SUFBC_2_D complex. This explains why suppression of the *SUFB* gene has a more rapid and pronounced effect on complex stability and plant phenotype than suppression of the *SUFC* gene does. The differential degradation rates of the SUFBC_2_D components provide a mechanistic explanation for the observed variations in senescence phenotypes between the *SUFB*- and *SUFC*-RNAi lines following inducer application.

### Increasing the SUFB content leads to increased SUFC and SUFD content

The aforementioned results and our own previous results demonstrate that the suppression of *SUFB* and *SUFC* transcription ultimately leads to decreases in SUFB, SUFC, and SUFD (Hu et al. 2017) (Fig. 2E). To investigate the impact of increased SUFB and SUFC on the levels of SUFBC_2_D subunits, the contents of these proteins in 5-week-old WT, *SUFB*-OX and *SUFC*-OX plants were analysed by immunoblotting (Fig. 3). The relative expression levels of the *SUFB*, *SUFC*, and *SUFD* genes in *SUFB*-OX and *SUFC*-OX were first separately assessed to confirm the successful overexpression of the relevant genes in the transgenic plant materials without significant changes to the expression of the other two genes (Supplementary Fig. S4). At the protein level, when SUFB was overaccumulated by overexpressing *SUFB*, the contents of both SUFC and SUFD increased compared with those in the WT (Fig. 3). Conversely, in *SUFC*-OX leaves, an obvious increase in SUFC itself occurred, but the contents of SUFB and SUFD did not obviously change (Fig. 3). To make the immunoblot results more intuitive, quantitative analysis of protein bands was also conducted (Fig. 3, with corresponding standard curves in Supplementary Fig. S5). To further confirm the reliability of these results, we first determined via qRT-PCR that only the *SUFB* gene was overexpressed in the *SUFB*-OX plants at 5‒8 weeks old, while the *SUFC* and *SUFD* genes maintained normal levels (Supplementary Fig. S6). Immunoblot analysis was performed on the contents of SUFB, SUFC, and SUFD in the leaves of 6-to 8-week-old plants (Fig. 1B, D, F, H and J), and the corresponding data are shown with scatter plots indicating the relationships between each pair of subunits (Fig. 4). The trends of the changes in SUFB, SUFC and SUFD in the *SUFB*-OX and *SUFC*-OX plants relative to those in the WT plants remained consistent with those in the 5-week-old plants. These results confirmed that an increase in the SUFB content further elevated the contents of SUFC and SUFD, with a more pronounced increase in SUFD. A similar phenomenon was also observed in 6-8-week-old plants, which indicated that the regulation of SUFC and SUFD contents by SUFB is not a specific phenomenon occurring at a particular plant growth stage but rather a consistent occurrence throughout leaf development. On the basis of these findings, we propose the following hypothesis: SUFB may recruit the other two subunits in the assembly of the SUFBC_2_D complex, whereas SUFC and SUFD do not possess this capability.

**Figure 3.**
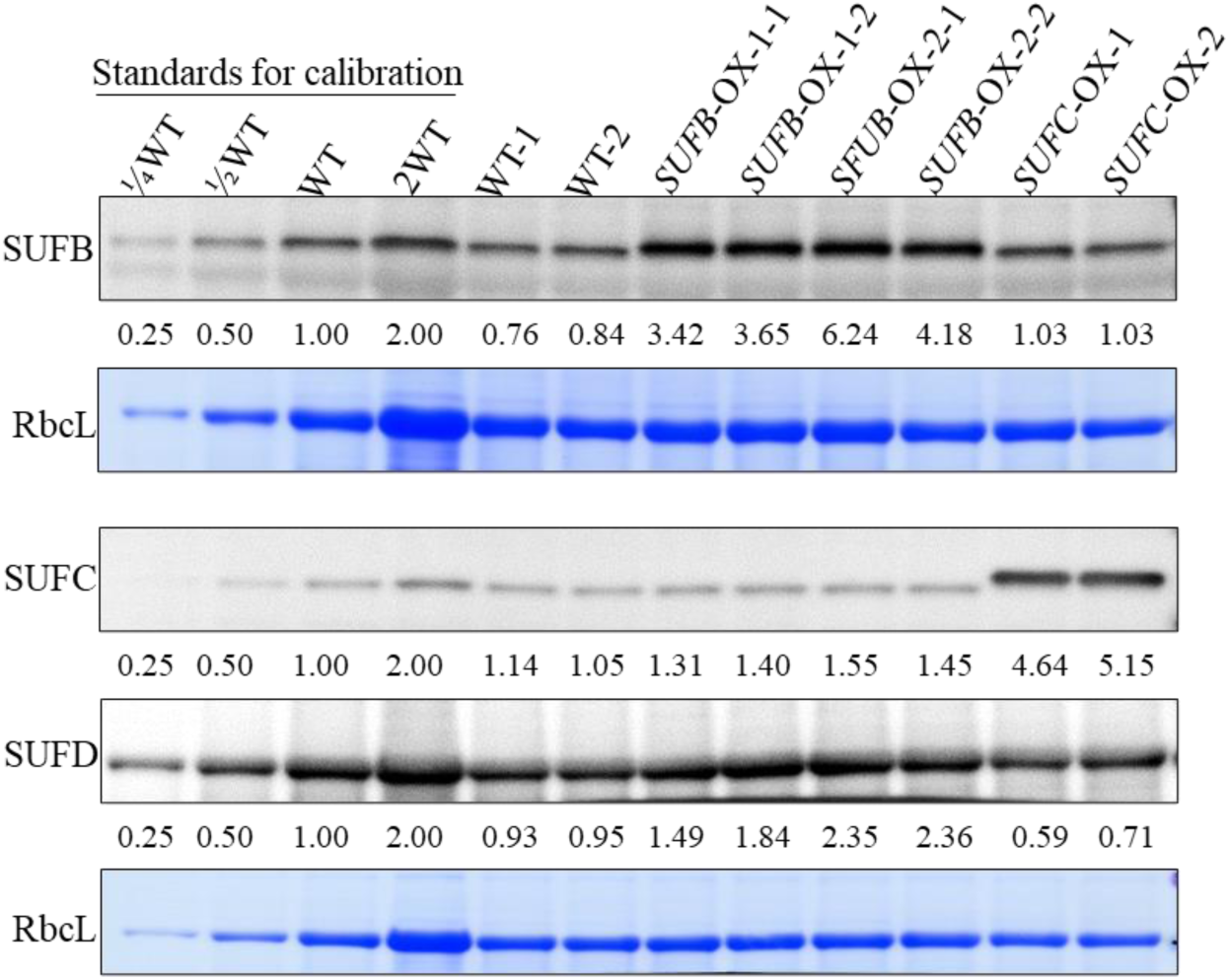
Immunoblot results showing the abundances of the SUFB, SUFC and SUFD proteins in the mature leaves of 5-week-old WT, *SUFB*-OX and *SUFC*-OX plants. The numbers at the bottom of each immunoblot indicate the relative quantities of the indicated proteins. CBB-stained bands of RbcL (Rubisco large subunit) are shown as loading controls. The major bands detected by α-SUFB in the *SUFB-OX* plants had higher molecular size because it is SUFB-FLAG other than SUFB. The major bands detected by α-SUFC in *SUFC-OX* plants had higher molecular size indicates SUFC-HA other than SUFC.

**Figure 4.**
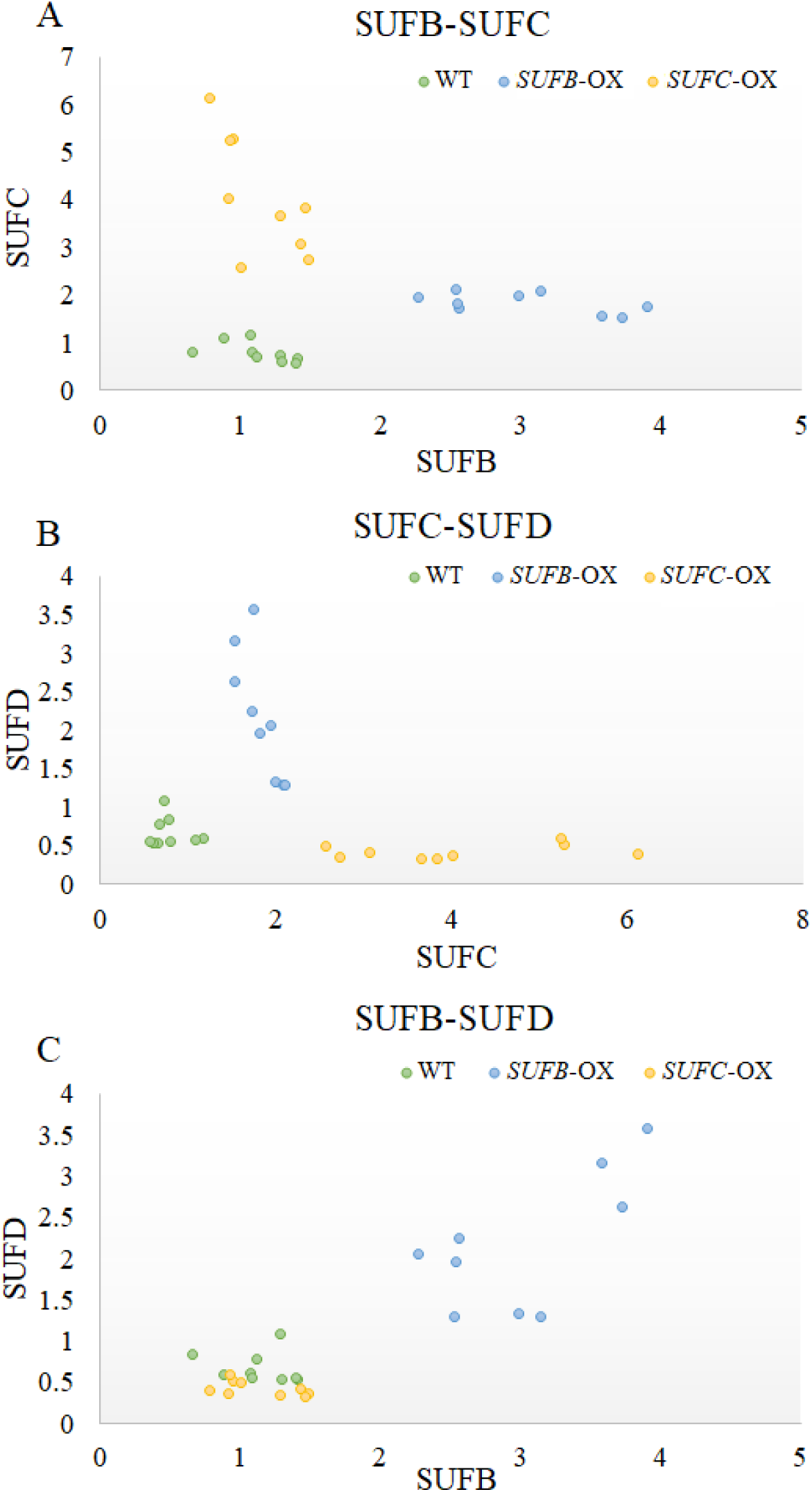
Content relationships among the three subunits of the SUFBC_2_D complex. (A) Relationship between SUFB and SUFC; (B) relationship between SUFC and SUFD; (C) relationship between SUFB and SUFD. The values on the x-axis and y-axis respectively represent the signal intensities corresponding to the immunoblot bands of the relevant proteins.

### SUFB regulates the content of SUFBC_2_D complexes

Since SUFB may stabilize SUFC and SUFD, we next sought to investigate whether the SUFB, SUFC, and SUFD proteins with increased abundance in the *SUFB*-OX lines were assembled into SUFBC_2_D. Two-dimensional (2D) BN/SDS‒PAGE was employed to analyse the state of complex formation in different lines. In the WT and *SUFB*-OX lines, both SUFB and SUFD were detected at approximately 170 kDa (larger than LHCII-T) and 110 kDa (between LHCII-T and LHC-M), and SUFC was also detected at approximately 170 kDa and at a low-molecular-mass position (between LHC-M and free pigments) (Fig. 5). Previous studies have shown that the 170 kDa spots correspond to the SUFBC_2_D complex (SUFB is approximately 60 kDa, SUFC is approximately 30 kDa, and SUFD is approximately 50 kDa), and the spots at 110 kDa are identified as the SUFBD complex(Hu et al. 2017). The spots at the low-molecular-mass positions correspond to SUFC. Therefore, in both the WT and *SUFB*-OX lines, SUFB and SUFD monomers are scarce, almost undetectable. Moreover, the proportion of monomeric SUFC to total SUFC in the *SUFB*-OX lines did not increase compared with that in the WT. These results suggest that the additional SUFC is at least partially assembled into the SUFBC_2_D complex, and that the additional SUFD is either entirely or predominantly assembled with SUFB into SUFBD or SUFBC_2_D. The proportions of monomeric SUFC to total SUFC in the *SUFC*-OX plants was significantly greater than that in the WT and *SUFB*-OX plants (Fig. 5). This result demonstrated that overexpressing SUFC did not increase the SUFBC_2_D content because there was no corresponding increase in SUFB or SUFD. Overall, we conclude that SUFB can stabilize SUFC and SUFD and promote SUFBC_2_D assembly, whereas SUFC does not have these capabilities.

**Figure 5.**
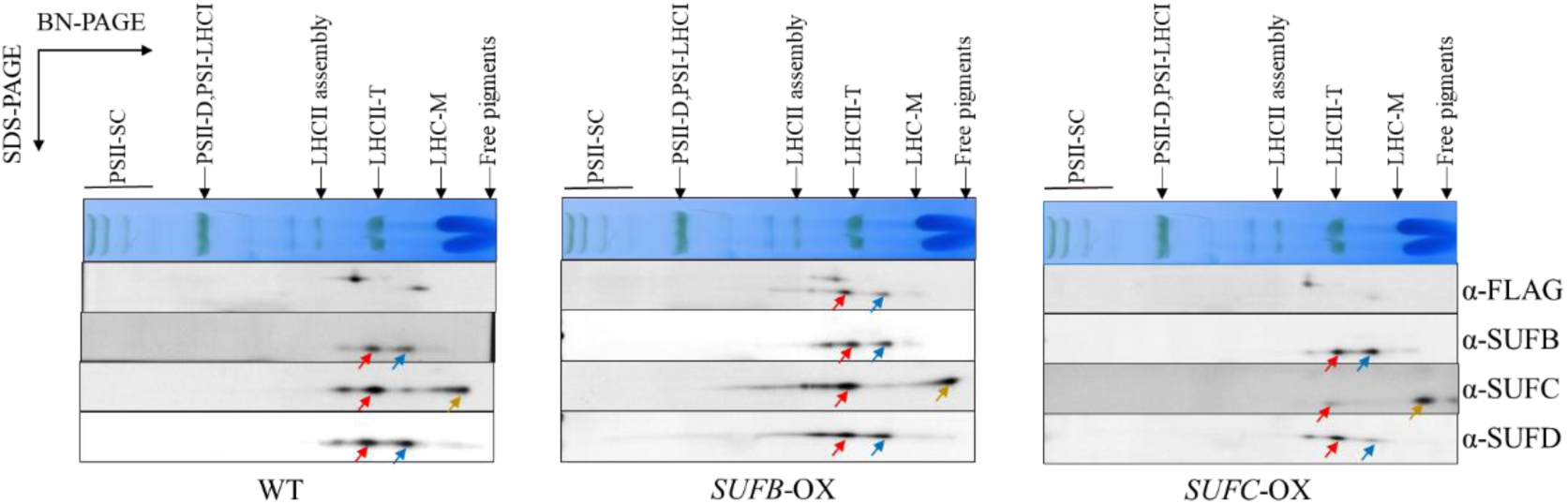
2D-BN/SDS‒PAGE analysis of the complexes that might contain SUFB, SUFC and/or SUFD, followed by immunoblotting. The primary antibodies used for immunoblotting were α-FLAG, α-SUFB, α-SUFC and α-SUFD. The deduced SUFBC_2_D complex is indicated with red arrows, and the molecular size is larger than that of the LHCII-T complex. The deduced SUFBD complex is indicated with blue arrows, and the molecular size is between the molecular sizes of the LHCII-T and LHC-M complexes. The SUFC signaling is indicated with yellow arrows, and the molecular size is close to that of free pigments.

### SUFB regulates the homeostasis of SUFBC_2_D in plants grown under iron deficiency conditions

The above studies revealed that artificially downregulated and upregulated SUFB contents in leaves decreased and increased the SUFBC_2_D content, respectively. We next examined whether SUFB regulates SUFBC_2_D content under specific conditions other than gene modification. It has been reported that the transcription of *SUFB* is downregulated at an early stage of iron deficiency, even before the downregulation of mRNAs encoding abundant chloroplastic ferritins (Kroh and Pilon 2020). Therefore, we tested whether the downregulation of *SUFB* transcription under iron-deficient conditions contributes to the modulation of the SUFBC_2_D content.

Uniform 10-day-old WT seedlings were transferred and grown on new half-strength Murashige and Skoog (½ MS) medium with or without iron supplementation for 3 days. The seedlings subjected to iron deficiency were relatively small, and the newly grown leaves were yellowish. After 6 days of iron deficiency, more young leaves presented yellowish phenotypes. DAB staining was used to examine H_2_O_2_ levels in 10-day-old WT plants with or without iron deficiency for 6 days, and the leaves of the plants with iron deficiency presented much greater H_2_O_2_ accumulation than those without iron deficiency (Supplementary Fig. S7). After 3-day-old iron-deficient seedlings were transferred to iron-containing ½ MS medium for 3 days, the yellow leaves returned to a green colour (Fig. 6A).

**Figure 6.**
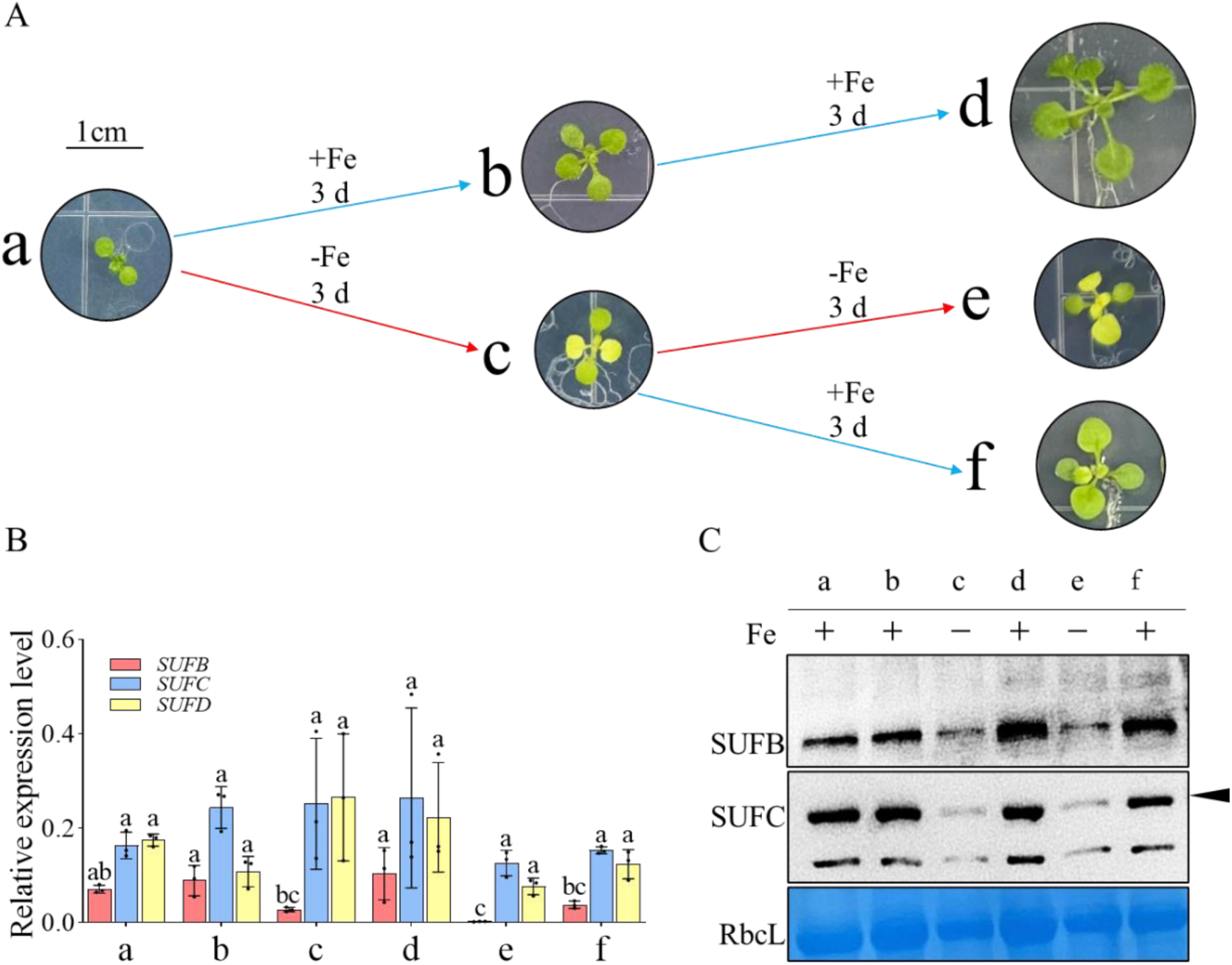
The levels of mRNA and protein in SUFs under Fe-sufficient (+Fe) and Fe-deficient (−Fe) conditions in WT plants. (A) WT plants were grown on +Fe and −Fe media containing 0.8% agar and 2% sucrose. a, 10-day-old WT grown on ½ MS medium (+Fe); b, 10-day-old WT grown on ½ medium (+Fe) for an additional 3 days; c, 10-day-old WT grown on −Fe medium for an additional 3 days; d, 10-day-old WT grown on +Fe medium for 6 days; e, 10-day-old WT grown on −Fe medium for 6 days; f, 10-day-old WT grown on −Fe medium for 3 days; thereafter, they were grown on +Fe medium for 3 days. Red arrows indicate seedlings that were grown under −Fe conditions; blue arrows indicate seedlings that were grown under +Fe conditions. Scale bar = 1 cm. (B) Relative expression levels of *SUFB*, *SUFC* and *SUFD* genes in WT plants grown under +Fe and −Fe conditions. *ACTIN2* was used as the internal control. The data are transcript levels relative to that of *ACTIN2* (with *ACTIN2* set to 1). The data represent the means ± SEs (n = 3 biological replicates). Different lowercase letters within the same coloured columns denote significant differences (*P* < 0.05), as determined by two-way ANOVA followed by Tukey’s post hoc test. (C) Immunoblot analysis of SUFB and SUFC contents in WT plants grown under +Fe and −Fe conditions. The black arrow indicates the SUFC bands. The smaller bands in the same blot may be the degradation products of SUFC. RbcL, Rubisco large subunit.

Measurement of the transcript levels of *SUFB, SUFC*, and *SUFD* revealed that *SUFB* responds to iron deficiency; its expression significantly decreased after 3 days of iron deficiency, and its mRNA level was nearly undetectable at the 6th day. The *SUFB* mRNA level was restored on the 3rd day after iron was provided to the iron-deficient seedlings. In contrast, the transcription of *SUFC* and *SUFD* remained relatively stable and did not significantly change with fluctuations in iron levels (Fig. 6B). We next examined the protein contents of SUFB and SUFC in seedlings under different iron conditions. The results revealed that the contents of SUFB and SUFC significantly decreased in the iron-deficient plants and increased when iron was supplied (Fig. 6C). These findings suggest that plants regulate the transcription level of *SUFB* in response to iron status and thereby regulate the content of SUFBC_2_D. On the basis of these results, we speculate that the reduction in the content of the SUFBC_2_D complex under iron deficiency conditions may be a strategy for plants to allocate iron to ensure their survival. To validate our hypothesis, we subjected 10-day-old WT, *SUFB*-OX, and *SUFC*-OX seedlings to iron deficiency treatment for 6 days. Compared with those of the WT and SUFC-OX lines, the newly grown leaves of both *SUFB*-OX lines presented more severe yellow phenotypes (Supplementary Fig. S8) and lower light-harvesting complex (LHC) content, which was consistent with the degree of the yellowing phenotype (Supplementary Fig. S9). The contents of SUFB and SUFC significantly decreased in both the WT and OX seedlings, whereas considerable amounts of SUFB and SUFC were present in the *SUFB*-OX and *SUFC*-OX lines, respectively. Artificially increasing the SUFB content in iron-deficient leaves resulted in more severe yellowing phenotypes, which suggests that the transcriptional regulation of *SUFB* plays an important role in the adaptation of leaves to iron deficiency. Because excess SUFB can assemble SUFBC_2_D, whereas SUFC cannot, we assume that the SUFBC_2_D complex needs to be decreased when iron deficiency occurs in plants.

### The SUFBC_2_D complex is degraded by the CLP system

The above studies suggest that SUFBC_2_D is quickly degraded when plants are subjected to iron deficiency. To determine which protease is involved in SUFBC_2_D degradation, we investigated the effects of major proteases on SUFBC_2_D. The two main protease systems in the chloroplast are the CLP system and the filamentation temperature-sensitive H (FTSH) system. Consequently, we ordered mutants in which these two proteases were impaired: *clpc1* and *var2-1* (*ftsh2* mutant) (Fig. 7A). The immunoblotting results revealed that SUFB, SUFC, and SUFD were all overaccumulated in *clpc1*, with contents several times greater than those in the WT. In contrast, the contents of these proteins in *var2-1* were similar to those in the WT (Fig. 7B). This finding suggests that the CLP system likely participates in the degradation of one or more subunits of SUFBC_2_D, thereby playing a regulatory role in SUFBC_2_D. Since the degraded substrates of the CLP system are recognized by the conserved substrate adaptor CLPS1 (Nishimura et al. 2013), protein-protein interaction assays were employed to examine whether any subunit of SUFBC_2_D can interact with CLPS1. First, AD-CLPS1, BD-SUFB, and BD-SUFC recombinant plasmids for the yeast two-hybrid (Y2H) assay were constructed, and the paired AD and BD recombinant plasmids were cotransformed into yeast cells. Subsequent analysis of the transformants confirmed the interaction between CLPS1 and SUFB/SUFC (Fig. 7C). In agreement with the Y2H results, the interaction between SUFB/SUFC and CLPS1 was further validated through a bimolecular fluorescence complementation (BiFC) assay in *Nicotiana benthamiana* (Fig. 7D). We prepared fusion constructs encoding the C-terminal fragment of the enhanced yellow fluorescent protein (cEYFP) fused to the full-length CDS of *SUFB* (*SUFB-cEYFP*) or *SUFC* (*SUFC-cEYFP*) under the control of the 35S promoter. cEYFP served as a negative control for CLPS1 interactions. We also created fusion constructs encoding the N-terminal fragment of the enhanced yellow fluorescent protein (*nEYFP*) fused to the CDS of *CLPS1* (*CLPS1-nEYFP*). nEYFP served as a negative control for SUFB or SUFC interaction. When *SUFB-cEYFP* and *SUFC-cEYFP* were transiently coexpressed with *CLPS1-nEYFP*, strong eYFP fluorescence was detected in the chloroplasts. No visible fluorescence was detected in the negative controls, in which *SUFB-cEYFP* and *SUFC-cEYFP* were coexpressed with *nEYFP*, and *cEYFP* was coexpressed with *CLPS1-nEYFP*(Fig. 7D). The BiFC results support interactions between SUFB/SUFC and CLPS1 *in vivo*.

**Figure 7.**
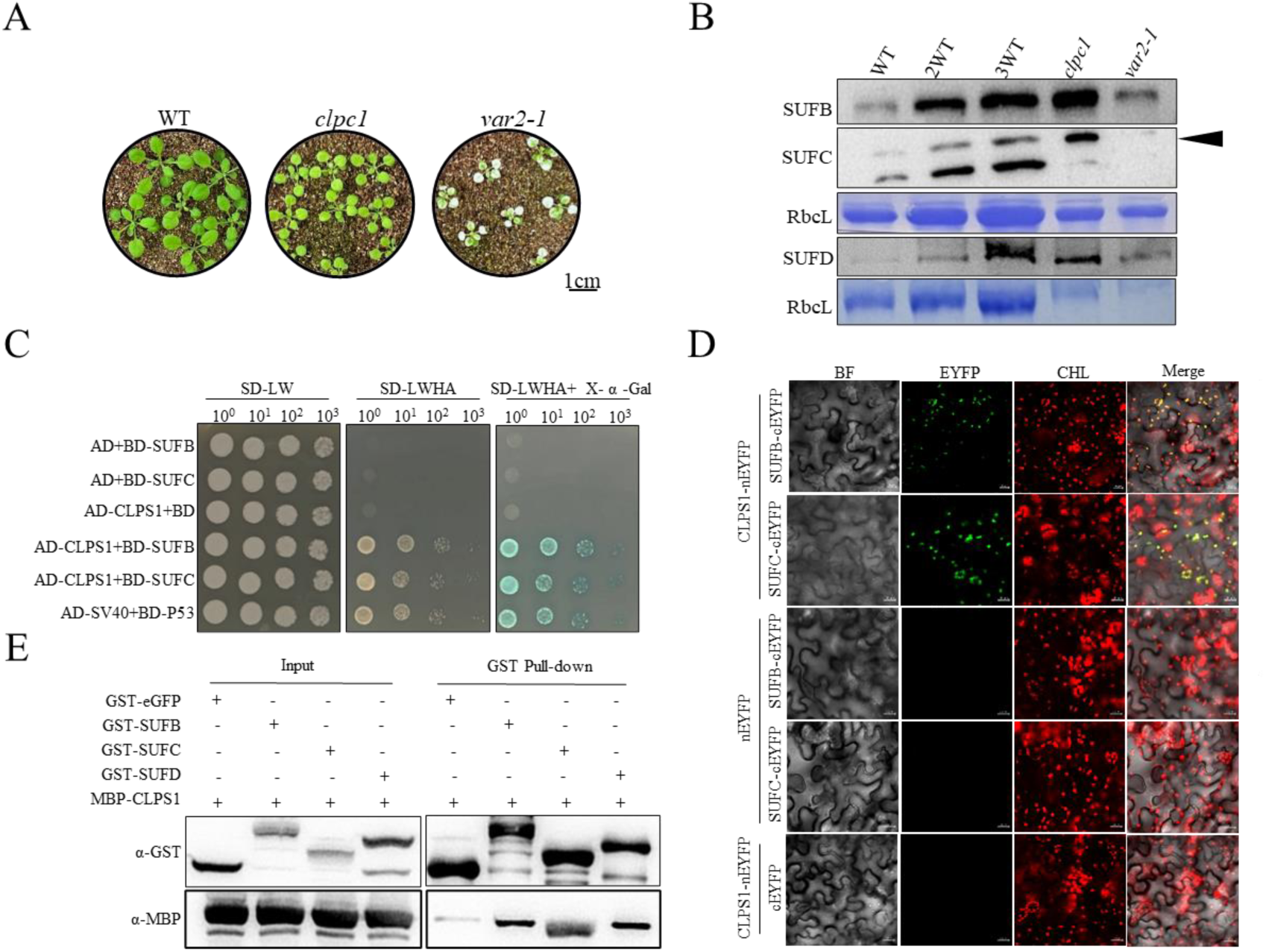
The chloroplast CLP system degrades SUFB, SUFC and SUFD. (A) Phenotypes of 3-week-old WT, *clpc1* and *var2-1* mutants grown in soil. Scale bar = 1 cm. (B) Immunoblot analysis of SUFB, SUFC and SUFD contents in WT, *clpc1* and *var2-1* mutants. The black arrow indicates the bands corresponding to SUFC. The smaller bands in the same blot may be the degradation products of SUFC. CBB-stained bands of the Rubisco large subunit (RbcL) are shown as loading controls. (C) Analyses of the interactions between CLPS1 and SUFB/SUFC via a yeast two-hybrid (Y2H) assay. The interactions between SUFB or SUFC fused with GAL4 BD and CLPS1 fused with GAL4 AD were examined. Positive control, the interaction between P53 fused with GAL4 BD and SV40 fused with GAL4 AD. Negative controls, the interaction between CLPS1 fused with GAL4 AD and GAL4 BD and the interaction between SUFB and SUFC fused with GAL4 BD and GAL4 AD. (D) BiFC analyses of the interactions between CLPS1 and SUFB/SUFC in the leaves of *N. benthamiana*. Scale bars = 20 µm. (E) Pull-down assay of the interactions between CLPS1 and chloroplast SUF components. GST-tagged SUFB, SUFC and SUFD were used as bait proteins. MBP-tagged CLPS1 was used as prey protein. The recombinant proteins were incubated with GST-tag purification resin and eluted, and analysis of the bound proteins was completed by immunoblotting.

The *in vitro* pull-down assay revealed that the anti-GST antibody pulled down recombinant MBP-CLPS1 in samples coincubated with GST-SUFB, GST-SUFC, and GST-SUFD but not with GST-eGFP (as a negative control) (Fig. 7E). Taken together, the results of three independent assays supported that SUFB/SUFC/SUFD all directly physically interact with CLPS1.

Overall, this study provides strong evidence that CLP protease regulates the protein levels of SUFB, SUFC and SUFD. The observed faster turnover of SUFB compared to other subunits, which is also critical for the regulation of SUFBC_2_D complex levels, can be attributed to CLP protease-mediated regulation.

## Discussion

The SUFBC_2_D complex is an Fe‒S cluster scaffold conserved in both bacteria and higher plants. The genes encoding these proteins are present in the *sufABCDSE* operon in bacteria; however, the expression of *SUFB*, *SUFC* and *SUFD* is not always synchronous in higher plants. First, the expression level of *SUFB* is greater than that of *SUFC* and *SUFD* in mature leaves (Liang et al. 2014) (Fig. 1A). Second, the transcription of *SUFB* was greatly increased when leaf senescence is induced by darkness, whereas the mRNA levels of *SUFC* and *SUFD* are high in green leaves (Nagane et al. 2010). Third, *SUFB* expression is downregulated early after the introduction of Fe deficiency, whereas *SUFC* and *SUFD* are less sensitive to Fe deficiency (Liang et al. 2014). In this study, we measured the transcript levels of *SUFB*, *SUFC*, and *SUFD* in the naturally aged leaves of 4-8-week-old WT plants via qRT‒PCR. The results indicated that only *SUFB* was induced during natural leaf senescence (Fig. 1A), which is consistent with the results of previous studies. However, at the protein level, the immunoblotting results revealed that the contents of SUFB, SUFC and SUFD did not significantly change during natural leaf senescence (Fig. 1B-K). In contrast, Nagane et al. reported that significant accumulation of SUFB protein occurs during dark-induced senescence in leaves (Nagane et al. 2010). We suggest two reasons for the inconsistent SUFB accumulation during dark-induced and natural leaf senescence. First, the transcript level of *SUFB* increased by 2.5 fold during dark-induced leaf senescence (Nagane et al. 2010), whereas it increased by approximately 2 fold during natural leaf senescence. Second, some chloroplastic Fe‒S proteins accumulate during leaf senescence, including HCAR and PAO, the enzymes that catalyse chlorophyll degradation (Hirashima et al. 2009; Meguro et al. 2011). The senescence process is accelerated by incubation in the dark; to provide enough Fe‒S clusters for these senescence-related apoproteins, SUFBC_2_D may therefore accumulate rapidly in leaves. The natural senescence of Arabidopsis leaves is a long process that lasts 20-40 days. Therefore, senescence requires that Fe‒S proteins accumulate much slower. It is necessary to maintain the content of SUFBC_2_D to provide sufficient Fe‒S clusters to these Fe‒S proteins. At the end of leaf senescence, SUFBC_2_D also gradually decreases with the completion of essential senescence processes such as chlorophyll degradation. At this time, fewer and fewer Fe‒S proteins are needed.

During leaf senescence, the transcription level of *SUFB* is notably increased, whereas the protein levels of SUFB, SUFC, and SUFD are not significantly increased. We propose that SUFB is highly susceptible to degradation during leaf senescence in Arabidopsis because of the senescence-specific increase in H_2_O_2_ (Bieker et al., 2012; Fig. 2B). In fact, SUFB is the major subunit that binds Fe‒S, which is sensitive to ROS (Sutton et al. 2004). The SUFBC_2_D complex is required to provide Fe‒S clusters for the biosynthesis of Fe‒S proteins in chloroplasts. To maintain the content of SUFBC_2_D, plants increase the transcription of *SUFB* to enable the rapid synthesis of SUFB, aiming to balance the accelerated rate of SUFB degradation during leaf senescence. To test this hypothesis, we first obtained previous transcriptome data showing that *SUFB* transcription is induced by oxidative stress and leaf senescence (Fig. 1A, Supplementary Fig. S1 and S3), which initially supports our hypothesis. We further found that when *SUFB*-RNAi was induced, mature green leaves could not undergo senescence programmatically and died before turning yellow (Fig. 2A). In *SUFC*- and *SUFD*-RNAi plants, the mature leaves of the plants presented normal senescence phenotypes, although RNAi was also induced (Fig. 2A). The different senescence phenotypes of these lines were caused by the different SUFBC_2_D contents in their leaves. The rates of decrease in the SUFB and SUFC contents in the mature leaves of the *SUFB*-RNAi plants were significantly faster than those in the *SUFC*-RNAi plants after they were sprayed with the inducer Dex (Fig. 2E). Because the transcription levels of *SUFB* and *SUFC* are repressed to similar levels in the two lines, respectively (Hu et al. 2017), these results indicated that the SUFB turnover rate is faster than the SUFC turnover rate. Although we did not determine the turnover rate of SUFD, considering the senescence phenotype of the *SUFD*-RNAi plants after Dex treatment (Fig. 2A), we infer that SUFD has a slower turnover rate than SUFB. Interestingly, the results also suggest that if the SUFB or SUFC content is insufficient, the other unassembled protein is also degraded quickly. Taken together, SUFB is the limiting factor for the stability of the SUFBC_2_D complex, because SUFB is the subunit with the fastest turnover rate.

To further investigate whether SUFB can positively regulate the SUFBC_2_D complex, *SUFB*-OX and *SUFC*-OX plants were analysed by immunoblotting. The results revealed that only the increase in SUFB protein levels further increased the content of the other two subunits of the SUFBC_2_D complex, suggesting that SUFB may have the ability to stabilize SUFC and SUFD (Fig. 1 and 3). In contrast, although the SUFC content was significantly increased in the *SUFC*-OX line, the contents of SUFB and SUFD were similar to those in the WT. Moreover, a similar trend was observed when the plants were under iron deficiency stress conditions (Supplementary Fig. S9). To further confirm whether the additional SUFB, SUFC, and SUFD form complexes, 2D BN/SDS‒PAGE was employed, and the results revealed that, in *SUFB*-OX plants, the increased SUFB, SUFC, and SUFD assembleed into SUFBC_2_D or SUFBD, whereas in *SUFC*-OX plants, only the proportions of SUFC monomers significantly increased. The excess SUFC did not form SUFBC_2_D (Fig. 5). We infer that SUFB is the core component of the SUFBC_2_D complex, whereas SUFC and SUFD play supporting roles during Fe‒S cluster assembly. This hypothesis is supported by research in *E. coli*, which suggested that SufB is the scaffold protein of the Suf system. It can transiently assemble an Fe‒S cluster *in vitro* (Layer et al. 2007; Blahut et al. 2020). SufC can increase the interaction between SufB and SufE to promote the transfer of sulfur from SufE to SufB. SufD and SufC also assist in SUFBC_2_D iron acquisition *in vivo* (Blahut et al. 2020). The mechanisms of the different functions in stabilizing the SUFBC_2_D complex between SUFB and SUFC require further investigation.

Iron is the fourth most abundant element in the Earth’s crust, but it is not readily available for biological use. Iron in soil exists mainly in the form of iron oxides, which are not very readily available to plants (Colombo et al. 2014; Connorton et al. 2017). Therefore, iron deficiency is a common stress in plants. The transcript level of *SUFB* is downregulated in the early stage of iron deficiency (Xu et al. 2005; Kroh and Pilon 2020), and this effect is highly conserved from plants (Liang et al. 2014; Pan et al. 2015; Hantzis et al. 2018) to cyanobacteria (Georg et al. 2017), whereas the expression of other *SUF* proteins (*SUFA*, *SUFC*, *SUFD*, *SUFE*, and *SUFS*) shows no significant differences with or without iron deficiency (Liang et al. 2014; Kroh and Pilon 2020). In contrast, iron stimulates not only the ATPase activity of SUFB but also *SUFB* expression at the transcriptional level (Xu et al. 2005). These findings revealed a novel characteristic of *SUFB* in respond to iron levels in plants.

At the protein level, our findings revealed that SUFB rapidly decreases in response to iron deficiency conditions (Fig. 6C), resulting in a significant reduction in the content of the SUFBC_2_D complex. We hypothesize that degradation of the SUFBC_2_D complex is necessary for plant survival under iron deficiency conditions. *In vivo*, in addition to Fe‒S clusters, Fe exists in other forms such as nonhaem iron, haem and siroheme (Kroh and Pilon 2020). These forms of Fe function as cofactors of proteins that could also be crucial for plant survival. For example, *Xantha-I* encodes the membrane subunit of the oxygen-dependent MPE (Mg-protoporphyrin IX monomethyl ester) cyclase, a key enzyme involved in chlorophyll biosynthesis. Mutants that are completely blocked at the *Xantha-I* locus are unable to produce chlorophyll (Rzeznicka et al. 2005). Therefore, we believe that plants need to reduce their SUFBC_2_D content under iron deficiency conditions, possibly to allocate Fe to other forms to maintain survival. This hypothesis was initially supported by the increased severity of the yellowing phenotype of *SUFB*-overexpressing lines grown under iron deficiency conditions (Supplementary Fig. S8 and S9). Further investigation is needed to demonstrate this possibility.

Chloroplast proteolysis is a key process in maintaining chloroplast protein homeostasis, ensuring optimal levels of functional proteins and removing aggregated, misfolded or unwanted proteins. There are multiple protease systems in chloroplasts, among which two energy-dependent, bacterial-derived proteases, FTSH2 and CLP, have been well studied. FTSH2s are membrane-embedded proteolytic complexes that contain an AAA (ATPase related to various cellular activities) and a Zn^2+^ metalloprotease domain. Chloroplast FTSH2s play important roles in the assembly and maintenance of the chloroplast membrane system (Wagner et al. 2012). These proteins are mainly responsible for the degradation of the photodamaged D1 protein of the photosystem II reaction centre (Kato et al. 2009; Sun et al. 2023), and unassembled thylakoid proteins (Ostersetzer and Adam 1997; Malnoë et al. 2014). The ATP-dependent serine-type CLP protease system is located in the chloroplast stroma and predominantly targets soluble proteins in the stroma, including protein degradation products from the thylakoid or envelope (Yuan and Van Wijk 2024). The SUFBC_2_D complex is the soluble protein complex located in the chloroplast stroma (Hu et al. 2017). Our results revealed that SUFB, SUFC, and SUFD were overaccumulated in *clpc1*, whereas the contents of these proteins in *var2-1* were similar to those in WT (Fig. 7B). Proteome analysis revealed that SUFC protein was overaccumulated in the tobacco *clpc1* mutant (Moreno et al. 2018). These results suggested that SUFB, SUFC, and SUFD may be degraded by the CLP protease system. The substrate proteins must be recognized by the recognition protein CLPS1 in the CLP system to be further degraded (Nishimura et al. 2013). For example, CLPS1 affinity studies revealed several chloroplast stromal targets that interact with CLPS1. In this study, Y2H and BiFC assays were employed to confirm that SUFB and SUFC directly interact with CLPS1, respectively (Fig. 7C and D). In addition, the results of the pull-down assay further demonstrated that SUFB, SUFC and SUFD all interact with CLPS1 (Fig. 7E). These results indicate that SUFB, SUFC and SUFD are all degradation substrates of the CLP system.

To summarize, we propose that plants upregulate and downregulate the transcriptional level of *SUFB* to regulate the content of SUFBC_2_D, thereby responding promptly to environmental changes. At the posttranslational level, the CLP system is responsible for degrading SUFB, SUFC and SUFD. Owing to the shorter half-life time of SUFB than that of SUFC (and possibly SUFD), the CLP system primarily regulates the levels of SUFBC_2_D by degrading SUFB firstly (Fig. 8).

**Figure 8.**
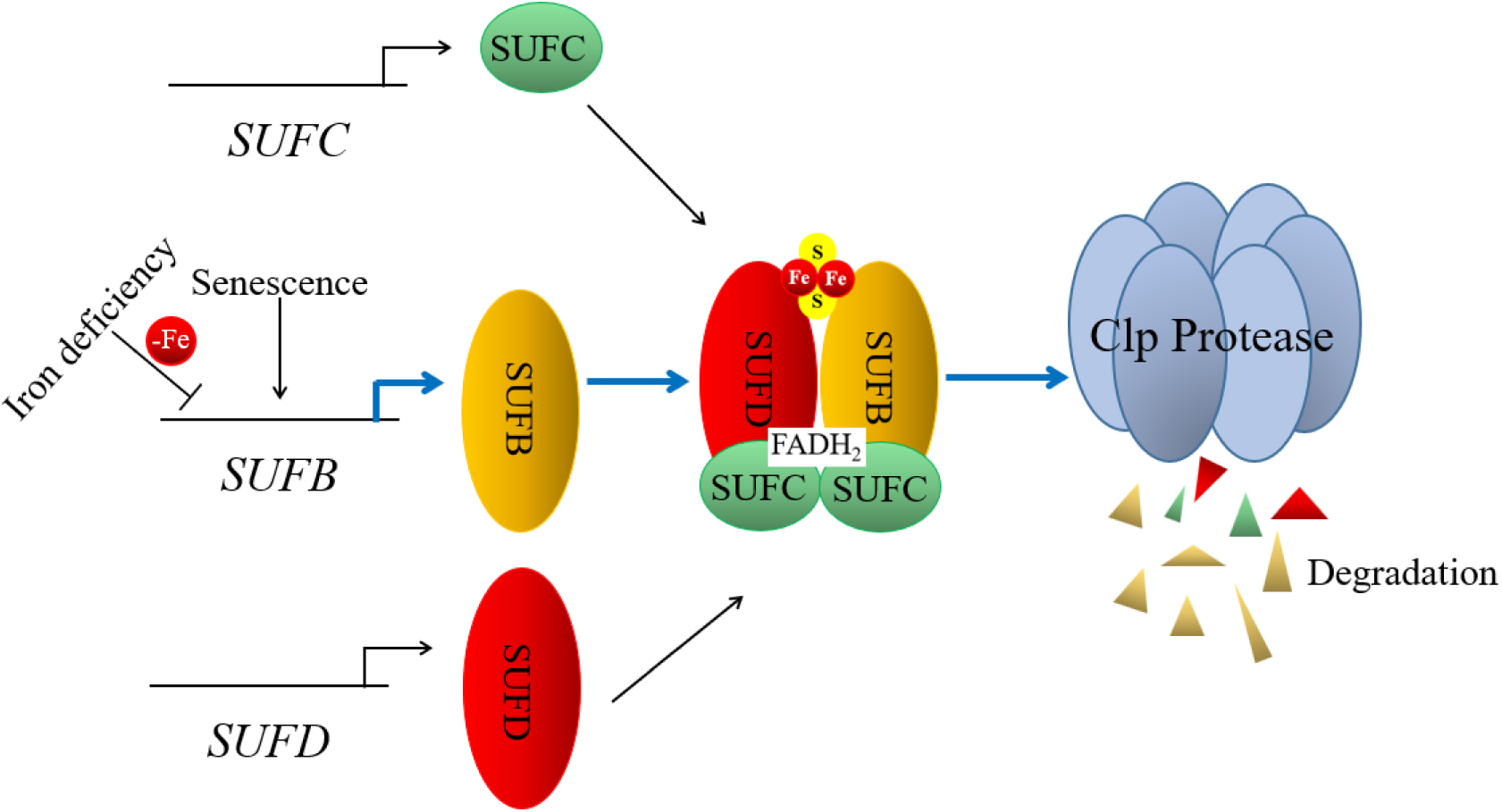
A model describing the mechanism by which plants manipulate *SUFB* to regulate the SUFBC_2_D complex in response to leaf senescence and iron deficiency. Under iron deficiency conditions, the transcript level of *SUFB* rapidly decreases, whereas during leaf senescence, it increases, resulting in the corresponding synthesis of less and more SUFB protein, respectively. This occurred because SUFB, SUFC and SUFD are all degraded by CLP protease, and SUFB has a shorter half-life than the other two components do. A decreased rate of SUFB synthesis resulted in less retention of the SUFBC_2_D complex under iron deficiency conditions. During leaf senescence, although the biosynthesis rate of SUFB increased, the degradation rate of SUFB also increased owing to the damage to SUFB caused by increased ROS during leaf senescence. Thus, the synthesis and degradation rates of SUFB were relatively balanced during the early stage of leaf senescence, maintaining the content of the SUFBC_2_D complex.

## Materials and methods

### Plant materials and growth conditions

All experimental materials from *Arabidopsis thaliana* were of the ecotype *Col-0* background. *Nicotiana benthamiana* was used for transient expression and BiFC assays. The *SUFB*-overexpressing lines (*SUFB*-OX-1 and *SUFB*-OX-2), the *SUFC*-overexpressing line (*SUFC*-OX) and the Dex-inducible RNAi lines *SUFB*-RNAi, *SUFC*-RNAi, and *SUFD*-RNAi were all described previously (Hu et al. 2017). Arabidopsis T-DNA insertion mutants *clpc1* (SALK_200928C) and *var2-1* (CS272) were purchased from the Arabidopsis Biological Resource Center (ABRC).

To obtain seedlings grown on sterilized medium, all the Arabidopsis seeds were surface sterilized with 75% (v/v) ethanol for 10 min, followed by a second round of surface sterilization with a small amount of 95% (v/v) ethanol for 1 min. The sterilized seeds were subsequently transferred to sterilized paper and sown on ½ MS medium supplemented with 0.8% (w/v) agar and 2% (w/v) sucrose. For the Fe-deficiency treatments, Fe was omitted from ½ MS, and standard ½ MS was employed for the Fe-sufficient conditions. To obtain plants grown in soil, mature seeds were sown directly in peat soil. The medium or soil with seeds was subsequently stratified at 4°C for 3 d, and the seeds were then grown in a growth chamber with 80-100 µmol photons m^-2^ s^-1^ fluorescent light at 23°C under long-day conditions (16 h light/8 h dark cycle).

### RNA extraction and transcript analysis

Total RNA was extracted from Arabidopsis leaves via the RNeasy Plant Mini Kit (Qiagen, Germany) according to the manufacturer’s instructions and reverse transcribed into cDNA using the PrimeScript RT reagent kit (Takara, Japan). SsoAdvanced Universal SYBR Green Supermix was used for qRT‒PCR (Bio-Rad, USA). The relative expression of selected genes was normalized to the expression of the *ACTIN2* (*AT3G18780*) gene, which was used as an internal control (Garapati et al. 2015; Cui et al. 2024). The qRT‒PCR primers used for analysis are listed in Supplementary Table S1.

### Sodium dodecyl sulfate (SDS) –PAGE and immunoblot analysis

Total protein was extracted from Arabidopsis leaves using 10 volumes (v/w) of protein extraction buffer containing 5% (v/v) 1 M Tris-HCl (pH 8.0), 2% (w/v) dodecyl lithium sulfate, 12% (w/v) sucrose and 1.5% (w/v) dithiothreitol. Proteins were separated on 14% polyacrylamide gels and subsequently transferred onto PVDF membranes. Specific primary antibodies against SUFB, SUFC and SUFD were produced in rabbits against recombinant Arabidopsis SUFB, SUFC and SUFD, respectively, expressed in *E. coli* (Hu et al. 2017). Anti-FLAG antibody was purchased from Agrisera (Vännäs, Sweden), and anti-GST and anti-MBP antibodies were purchased from ABclonal (Wuhan, China). Horseradish peroxidase (HRP)-labelled secondary antibodies were used, and HRP activity was detected via the ECL Plus western blotting detection system (PerkinElmer, USA) according to the manufacturer’s protocol.

### Relative quantitative analysis of the abundance of target proteins

Relative target protein quantification was conducted by utilizing a dilution series of the WT total protein as a control. The graphs were generated with the GraphPad Prism 9 software, and the signal intensity values were determined via CSAnalyzer4 software.

### *SUFB, SUFC* and *SUFD* RNAi induction

Dex was first dissolved in dimethyl sulfoxide (DMSO) to a concentration of 20 mM, and then diluted with 0.02% Tween-20 aqueous solution to give a final concentration of 20 µM Dex. To induce RNAi silencing, 3- or 4-week-old (dependent on different experiments) plants grown in soil were sprayed with 20 µM Dex solution.

### DAB staining

The detached leaves were harvested and stained with a DAB staining kit (Coolaber, Beijing, China) according to the manufacturer’s instructions. The leaves were subsequently observed and imaged under a stereomicroscope.

### Chlorophyll fluorescence measurements

The plants to be tested were first dark adapted for 30 min, and the maximal efficiency of PSII (*F*_v_*/F*_m_) was then determined via MultispeQ V2.0 (https://www.photosynq.com).

### Ion leakage

The 7^th^ and 8^th^ (leaves were numbered from the bottom) leaves of the plants were soaked in 2 mL of distilled water in centrifuge tubes and were shaken for 2 h, and the electrolyte leakage of the leaves was measured with a compact conductivity meter (LAQUAtwin-EC-33; http://www.horiba.com/jp). The leaves were subsequently boiled for 15 min and shaken for 2 h. The conductivity was measured again to assess the total electrolyte leakage from the leaves.

### Two-dimensional (2D)-blue native (BN)/SDS–PAGE

BN–PAGE was performed according to previously described methods (Wittig et al. 2006). Purified thylakoid membranes of the WT as protein marker of molecular size were prepared according to Salvi et al. (Salvi et al. 2008). Briefly, the purified thylakoid membrane proteins (which corresponded to 100 µg of Chl) were suspended in 100 µL of ice-cold BN solubilization buffer containing 50 mM imidazole-HCl (pH 7.0), 10% glycerol, 0.5 M 6-aminocaproic acid and 1 mM EDTA [with 1% protease inhibitor cocktail (Sigma)] and then solubilized with 1% (w/v) α-dodecyl maltoside (α-DM) on ice for 5 min. After centrifugation at 21,500 ×*g* for 2 min at 4°C, the supernatants were separated on 4–13% polyacrylamide gradient gels.

The samples for complex analysis were prepared by the following procedure. Leaves were homogenized in solubilization buffer and centrifuged for 10 min at 21,500 ×*g* at 4°C. The supernatants were collected and mixed with an equal volume of 2% (w/v) α-DM solution, and the proteins were separated on 4–13% polyacrylamide gradient gels at 4°C [anode buffer: 25 mM imidazole-HCl (pH 7.0, 4°C); cathode buffers: 50 mM tricine, 7.5 mM imidazole-HCl (pH 7.0, 4°C) and 0.02% Coomassie brilliant blue (CBB, G-250)] according to the methods described by Wittig et al.(Wittig et al. 2006).

Proteins in the BN–PAGE gel strips were denatured in solution buffer [1% (w/v) SDS and 50 mM DTT] for 30 min at room temperature and separated on a 2D–SDS– PAGE gel (14% acrylamide gel containing 6 M urea) via the Laemmli buffer system (Laemmli 1970). Immunoblotting was performed as described above.

### Y2H assay

The coding sequence (CDS) of *CLPS1* was cloned and inserted into the prey vector pGADT7, and the CDSs of *SUFB* and *SUFC* were cloned and inserted into the bait vector pGBKT7. The prey and bait plasmids were subsequently transformed into yeast strain Y187 or Y2HGold, respectively, and subsequently mated. The yeast cells were cultured on synthetic defined (SD) medium without -Leu-Trp. Interactions between the bait and prey proteins were further examined on SD medium (X-α-Gal and aureobasidin A) without -Leu-Trp-His-Ade. The interaction between the pGADT7-SV40 and pGBKT7-P53 proteins was used as a positive control. The primers used for the construction of Y2H vectors are listed in Supplementary Table S1.

### BiFC assay

BiFC assays were conducted following a previously described method (Kerppola 2008). In this experiment, the complete CDSs of *SUFB*, *SUFC* and *CLPS1* were amplified and recombined into pSAT4A-cEYFP-N1 and pSAT4A-nEYFP-N1 plasmids, respectively. These constructs and the empty vectors were separately transformed into *Agrobacterium tumefaciens* GV3101 (pSoup). The transformed *Agrobacteriums* cells were combined into different pairs, which were coinfiltrated into the leaves of *N. benthamiana* plants. The fluorescence signals were observed 2 day later via an LSM880NLO confocal microscope (Carl Zeiss, Germany). The primers used for the construction of the BiFC are listed in Supplementary Table S1.

### *In vitro* pull-down assay

The CDSs of the bait proteins (SUFB, SUFC and SUFD) were cloned and inserted into pGEX-4T-1 expression vectors to generate fusion constructs with glutathione S-transferase (GST) tags. The CDS of the prey protein (CLPS1) was cloned and inserted into the pET His6 MBP TEV LIC cloning vector (1 M) (#29656, Addgene) cloning vector to generate fusion constructs with an MBP tag. These constructs were subsequently transformed into the *E. coli* strain BL21 (DE3). The genes were expressed in *E. coli* in the presence of isopropyl β-D-1-thiogalactopyranoside (IPTG). For pull-down assays, protein extracts containing GST-fused bait proteins were incubated with GST-tag purification resin for 2–3 h at 4°C. After incubation, the resins were collected via simple centrifugation and then washed three times with PBS buffer [0.8% (w/v) NaCl, 0.02% (w/v) KCl, 0.4% (w/v) Na_2_HPO_4_·12H_2_O, 0.03% (w/v) KH_2_PO_4_, 0.02% (w/v) DTT and 0.02% (w/v) PMSF]. The protein extracts containing MBP-fused prey proteins were subsequently added and incubated with pull-down buffer [5% (v/v) 1 M Tris-HCl (pH 8.0), 6.7% (v/v) 3 M NaCl, 1% (v/v) 1 M MgCl_2_·6H_2_O, 0.2% (v/v) 0.5 M EDTA, 0.02% (w/v) DTT, 0.02% (w/v) PMSF and 1% (v/v) NP-40] for 3 h at 4°C. After incubation, the resins were collected via simple centrifugation and then washed 5 times with pull-down buffer. Finally, the precipitated proteins were eluted in 2 × SDS loading buffer [25% (v/v) 1 M Tris-HCl (pH 6.8), 10% (w/v) SDS, 41.67% (v/v) glycerol, 10% (v/v) 5 M DTT and 0.5% (w/v) bromophenol blue (BPB)], heated at 100°C for 5 min, and subjected to SDS–PAGE. Immunoblotting was used to detect the targeted recombinant proteins with commercial mouse anti-GST and anti-MBP antibodies, which were purchased from ABclonal (Wuhan, China). The primers used for the construction of the pull-down vectors are listed in Supplementary Table S1.

### Statistical analysis

Following confirmation of normality and homogeneity of variance, Tukey’s post hoc test was used for significant difference analysis with one-way ANOVA and two-way ANOVA among multiple samples, at a 95% confidence interval. All values are presented with mean ± s.d.as indicated. Data points are plotted onto the graphs, and the number of samples is indicated in the corresponding figure legends. P_value_ < 0.05 was considered to indicate statistical significance. The statistical analyses were performed using IBM SPSS statistics, version 29.0.2.0.

## Acknowledgements

We thank Junko Kishimoto at Hokkaido University for technical assistance in the laboratory. This work was funded by the National Natural Science Foundation of China (No. 32000197 and No. 31870265 to X.H.)

## Author contributions

Yuting Cheng, Conceptualization, Formal analysis, Investigation, Visualization, Methodology, Writing – original draft, Writing – review and editing; Zhaoyang Liu, Bing Yang, Investigation; Qingsong Jiao, Hisashi Ito, Atsushi Takabayashi, Conceptualization, Supervision, Ryouichi Tanaka, Ting Jia, Conceptualization, Supervision, Funding acquisition, Project administration, Writing – review and editing. Xueyun Hu, Conceptualization, Supervision, Funding acquisition, Project administration, Writing – original draft, Writing – review and editing.

## Reference

1. Balk J and Lobréaux S. Biogenesis of iron–sulfur proteins in plants. Trends Plant Sci. 2005, 10(7): 324–331. 10.1016/j.tplants.2005.05.002

2. Bieker S, Riester L, Stahl M, Franzaring J, and Zentgraf U. Senescence-specific alteration of hydrogen peroxide levels in *Arabidopsis thaliana* and oilseed rape spring variety *Brassica napus* L. cv. Mozart^F^. J Integr Plant Biol. 2012, 54(8): 540–554. 10.1111/j.1744-7909.2012.01147.x

3. Blahut M, Sanchez E, Fisher CE, and Outten FW. Fe-S cluster biogenesis by the bacterial Suf pathway. Biochim Biophys Acta Mol Cell Res. 2020, 1867(11): 118829. 10.1016/j.bbamcr.2020.118829

4. Colombo C, Palumbo G, He J-Z, Pinton R, and Cesco S. Review on iron availability in soil: interaction of Fe minerals, plants, and microbes. J. Soils Sediments. 2014, 14: 538–548.

5. Connorton JM, Balk J, and Rodríguez-Celma J. Iron homeostasis in plants – a brief overview. Metallomics. 2017, 9(7): 813–823. 10.1039/C7MT00136C

6. Cui X, Fan X, Xu S, Wang S, Niu F, Zhao P, Yang B, Liu W, Guo X, and Jiang Y-Q. WRKY47 transcription factor modulates leaf senescence through regulating PCD-associated genes in Arabidopsis. Plant Physiol Biochem. 2024, 213: 108805. 10.1016/j.plaphy.2024.108805

7. Garapati P, Xue G-P, Munné-Bosch S, and Balazadeh S. Transcription factor ATAF1 in Arabidopsis promotes senescence by direct regulation of key chloroplast maintenance and senescence transcriptional cascades. Plant Physiol. 2015, 168(3): 1122–1139. 10.1104/pp.15.00567

8. Garcia PS, D’Angelo F, Ollagnier De Choudens S, Dussouchaud M, Bouveret E, Gribaldo S, and Barras F. An early origin of iron–sulfur cluster biosynthesis machineries before Earth oxygenation. Nat Ecol Evol. 2022, 6(10): 1564–1572. 10.1038/s41559-022-01857-1

9. Georg J, Kostova G, Vuorijoki L, Schön V, Kadowaki T, Huokko T, Baumgartner D, Müller M, Klähn S, Allahverdiyeva Y, et al. Acclimation of oxygenic photosynthesis to iron starvation is controlled by the sRNA IsaR1. Curr Biol. 2017, 27(10): 1425–1436. 10.1016/j.cub.2017.04.010

10. Hantzis LJ, Kroh GE, Jahn CE, Cantrell M, Peers G, Pilon M, and Ravet K. A program for iron economy during deficiency targets specific Fe proteins. Plant Physiol. 2018, 176(1): 596–610. 10.1104/pp.17.01497

11. Hirashima M, Tanaka R, and Tanaka A. Light-independent cell death induced by accumulation of pheophorbide *a* in *Arabidopsis thaliana*. Plant Cell Physiol. 2009, 50(4): 719–729. 10.1093/pcp/pcp035

12. Hu X, Kato Y, Sumida A, Tanaka A, and Tanaka R. The SUFBC_2_D complex is required for the biogenesis of all major classes of plastid Fe-S proteins. Plant J. 2017, 90(2): 235–248. 10.1111/tpj.13483

13. Imlay JA. Iron-sulphur clusters and the problem with oxygen. Mol Microbiol. 2006, 59(4): 1073– 1082. 10.1111/j.1365-2958.2006.05028.x

14. Kato Y, Miura E, Ido K, Ifuku K, and Sakamoto W. The variegated mutants lacking chloroplastic FtsHs are defective in D1 degradation and accumulate reactive oxygen species. Plant Physiol. 2009, 151(4): 1790–1801. 10.1104/pp.109.146589

15. Kerppola TK. Bimolecular fluorescence complementation (BiFC) analysis as a probe of protein interactions in living cells. Annu Rev Biophys. 2008, 37: 465–487. 10.1146/annurev.biophys.37.032807.125842

16. Kroh and Pilon. Iron deficiency and the loss of chloroplast iron–sulfur cluster assembly trigger distinct transcriptome changes in Arabidopsis rosettes. Metallomics. 2020, 12(11): 1748– 1764. 10.1039/d0mt00175a

17. Laemmli UK. Cleavage of structural proteins during the assembly of the head of bacteriophage T4. Nature. 1970, 227(5259): 680–685. 10.1038/227680a0

18. Layer G, Gaddam SA, Ayala-Castro CN, Ollagnier-de Choudens S, Lascoux D, Fontecave M, and Outten FW. SufE transfers sulfur from SufS to SufB for iron-sulfur cluster assembly. J Biol Chem. 2007, 282(18): 13342–13350. 10.1074/jbc.M608555200

19. Lee K-C, Yeo W-S, and Roe J-H. Oxidant-responsive induction of the *suf* operon, encoding a Fe-S assembly system, through Fur and IscR in *Escherichia coli*. J Bacteriol. 2008, 190(24): 8244–8247. 10.1128/jb.01161-08

20. Liang X, Qin L, Liu P, Wang M, and Ye H. Genes for iron–sulphur cluster assembly are targets of abiotic stress in rice, *Oryza sativa*. Plant Cell Environ. 2014, 37(3): 780–794. 10.1111/pce.12198

21. Lill R. Function and biogenesis of iron–sulphur proteins. Nature. 2009, 460(7257): 831–838. 10.1038/nature08301

22. Lu Y. Assembly and transfer of iron–sulfur clusters in the plastid. Front Plant Sci. 2018, 9: 336. 10.3389/fpls.2018.00336

23. Malnoë A, Wang F, Girard-Bascou J, Wollman F-A, and De Vitry C. Thylakoid FtsH protease contributes to photosystem II and cytochrome *b*_6_*f* remodeling in *Chlamydomonas reinhardtii* under stress conditions. Plant Cell. 2014, 26(1): 373–390. 10.1105/tpc.113.120113

24. Meguro M, Ito H, Takabayashi A, Tanaka R, and Tanaka A. Identification of the 7-hydroxymethyl chlorophyll *a* reductase of the chlorophyll cycle in *Arabidopsis*. Plant Cell. 2011, 23(9): 3442–3453. 10.1105/tpc.111.089714

25. Mettert EL and Kiley PJ. Fe–S proteins that regulate gene expression. Biochim Biophys Acta Mol Cell Res. 2015, 1853(6): 1284–1293. 10.1016/j.bbamcr.2014.11.018

26. Moreno JC, Martínez-Jaime S, Schwartzmann J, Karcher D, Tillich M, Graf A, and Bock R. Temporal proteomics of inducible RNAi lines of Clp protease subunits identifies putative protease substrates. Plant Physiol. 2018, 176(2): 1485–1508. 10.1104/pp.17.01635

27. Nagane T, Tanaka A, and Tanaka R. Involvement of AtNAP1 in the regulation of chlorophyll degradation in *Arabidopsis thaliana*. Planta. 2010, 231(4): 939–949. 10.1007/s00425-010-1099-8

28. Nath K, Wessendorf RL, and Lu Y. A nitrogen-fixing subunit essential for accumulating 4Fe-4S-containing photosystem I core proteins. Plant Physiol. 2016, 172(4): 2459–2470. 10.1104/pp.16.01564

29. Nishimura K, Asakura Y, Friso G, Kim J, Oh S -h., Rutschow H, Ponnala L, and Van Wijk KJ. ClpS1 is a conserved substrate selector for the chloroplast Clp protease system in *Arabidopsis*. Plant Cell. 2013, 25(6): 2276–2301. 10.1105/tpc.113.112557

30. Ostersetzer O and Adam Z. Light-stimulated degradation of an unassembled Rieske FeS protein by a thylakoid-bound protease: the possible role of the FtsH protease. Plant Cell. 1997, 9(6): 957–965. 10.1105/tpc.9.6.957

31. Outten FW, Djaman O, and Storz G. A *suf* operon requirement for Fe–S cluster assembly during iron starvation in *Escherichia coli*. Mol Microbiol. 2004, 52(3): 861–872. 10.1111/j.1365-2958.2004.04025.x

32. Pan I-C, Tsai H-H, Cheng Y-T, Wen T-N, Buckhout TJ, and Schmidt W. Post-transcriptional coordination of the *Arabidopsis* iron deficiency response is partially dependent on the E3 ligases RING DOMAIN LIGASE1 (RGLG1) and RING DOMAIN LIGASE2 (RGLG2). Mol Cell Proteomics. 2015, 14(10): 2733–2752. 10.1074/mcp.M115.048520

33. Przybyla-Toscano J, Roland M, Gaymard F, Couturier J, and Rouhier N. Roles and maturation of iron–sulfur proteins in plastids. J Biol Inorg Chem. 2018, 23(4): 545–566. 10.1007/s00775-018-1532-1

34. Ravet K and Pilon M. Copper and iron homeostasis in plants: the challenges of oxidative stress. Antioxid Redox Signal. 2013, 19(9): 919–932. 10.1089/ars.2012.5084

35. Rzeznicka K, Walker CJ, Westergren T, Kannangara CG, Von Wettstein D, Merchant S, Gough SP, and Hansson M. *Xantha-l* encodes a membrane subunit of the aerobic Mg-protoporphyrin IX monomethyl ester cyclase involved in chlorophyll biosynthesis. Proc Natl Acad Sci. 2005, 102(16): 5886–5891. 10.1073/pnas.0501784102

36. Saini A, Mapolelo DT, Chahal HK, Johnson MK, and Outten FW. SufD and SufC ATPase activity are required for iron acquisition during in vivo Fe-S cluster formation on SufB. Biochemistry. 2010, 49(43): 9402–9412. 10.1021/bi1011546

37. Salvi D, Rolland N, Joyard J, and Ferro M. Purification and proteomic analysis of chloroplasts and their sub-organellar compartments. In. Organelle Proteomics, D Pflieger and J Rossier, eds. (Humana Press: Totowa, NJ), pp. 19–36. 10.1007/978-1-59745-028-7_2

38. Schmid M, Davison TS, Henz SR, Pape UJ, Demar M, Vingron M, Schölkopf B, Weigel D, and Lohmann JU. A gene expression map of *Arabidopsis thaliana* development. Nat Genet. 2005, 37(5): 501–506. 10.1038/ng1543

39. Sun Y, Li J, Zhang L, and Lin R. Regulation of chloroplast protein degradation. J Genet Genomics. 2023, 50(6): 375–384. 10.1016/j.jgg.2023.02.010

40. Sutton VR, Stubna A, Patschkowski T, Münck E, Beinert H, and Kiley PJ. Superoxide destroys the [2Fe-2S]^2+^ cluster of FNR from *Escherichia coli*. Biochemistry. 2004, 43(3): 791–798. 10.1021/bi0357053

41. Touraine B, Vignols F, Przybyla-Toscano J, Ischebeck T, Dhalleine T, Wu H-C, Magno C, Berger N, Couturier J, Dubos C, et al. Iron–sulfur protein NFU2 is required for branched-chain amino acid synthesis in Arabidopsis roots. J Exp Bot. 2019, 70(6): 1875– 1889. 10.1093/jxb/erz050

42. Wagner R, Aigner H, and Funk C. FtsH proteases located in the plant chloroplast. Physiol Plant. 2012, 145(1): 203–214. 10.1111/j.1399-3054.2011.01548.x

43. Winter D, Vinegar B, Nahal H, Ammar R, Wilson GV, and Provart NJ. An “electronic fluorescent pictograph” browser for exploring and analyzing large-scale biological data sets. PLoS ONE. 2007, 2(8): e718. 10.1371/journal.pone.0000718

44. Wittig I, Braun H-P, and Schägger H. Blue native PAGE. Nat Protoc. 2006, 1(1): 418–428. 10.1038/nprot.2006.62

45. Wollers S, Layer G, Garcia-Serres R, Signor L, Clemancey M, Latour J-M, Fontecave M, and Ollagnier De Choudens S. Iron-sulfur (Fe-S) cluster assembly. J Biol Chem. 2010, 285(30): 23331–23341. 10.1074/jbc.M110.127449

46. Xu XM, Adams S, Chua N-H, and Møller SG. AtNAP1 represents an atypical SufB protein in *Arabidopsis* plastids. J Biol Chem. 2005, 280(8): 6648–6654. 10.1074/jbc.M413082200

47. Yang B, Xu C, Cheng Y, Jia T, and Hu X. Research progress on the biosynthesis and delivery of iron–sulfur clusters in the plastid. Plant Cell Rep. 2023, 42(8): 1255–1264. 10.1007/s00299-023-03024-7

48. Ye H, Pilon M, and Pilon-Smits EAH. CpNifS-dependent iron-sulfur cluster biogenesis in chloroplasts. New Phytol. 2006, 171(2): 285–292. 10.1111/j.1469-8137.2006.01751.x

49. Yuan B and Van Wijk KJ. The chloroplast protease system degrades stromal DUF760-1 and DUF-2 domain-containing proteins at different rates. Plant Physiol. 2024, kiae431. 10.1093/plphys/kiae431

50. Zhang J, Bai Z, Ouyang M, Xu X, Xiong H, Wang Q, Grimm B, Rochaix J, and Zhang L. The DnaJ proteins DJA6 and DJA5 are essential for chloroplast iron–sulfur cluster biogenesis. EMBO J. 2021, 40(13): 106742. 10.15252/embj.2020106742

